# BioNE: Integration of network embeddings for supervised learning

**DOI:** 10.1101/2022.04.26.489560

**Authors:** Poorya Parvizi, Francisco Azuaje, Evropi Theodoratou, Saturnino Luz

## Abstract

A network embedding approach reduces the analysis complexity of large biological networks by converting them to lowdimensional vector representations (features/embeddings). These lower-dimensional vectors can then be used in machine learning prediction tasks with a wide range of applications in computational biology and bioinformatics. Several network embedding approaches have been proposed with different methods of generating vector representations. These network embedding approaches can be quite diverse in terms of data representation and implementation. Moreover, most were not originally developed for biological networks. Therefore comparing and assessing the performance of these diverse models in practice, in biological contexts, can be challenging. To facilitate such comparisons, we have developed the BioNE framework for integration of different embedding methods in prediction tasks. Using this framework one can easily assess, for instance, whether combined vector representations from multiple embedding methods offer complementary information with regards to the network features and thus better performance on prediction tasks. In this paper, we present the BioNE software suite for embedding integration, which applies network embedding methods following standardised network preparation steps, and integrates the vector representations achieved by these methods using three different techniques. BioNE enables selection of prediction models, oversampling methods, feature selection methods, cross-validation type and cross-validation parameters.

**Availability and implementation:** BioNE pipeline and detailed explanation of implementation is freely available on GitHub, at https://github.com/pooryaparvizi/BioNE

## Introduction

The advancement in high-throughput technologies has resulted in a substantial increase in available data, providing opportunities to research and gain a deeper understanding of interactions within biological systems. This allows for the analysis and exploration of cells or organisms as systems where molecular parts act together in a dynamic way (1). Network biology analysis, which is based on graph theory, could provide a structure for integrating these highthroughput multi-omics data and investigating interactions within biological systems (2). Although, analysing large amounts of data within the network is valuable, analysis and interpretation results can be challenging using conventional statistical methods (3). Network embedding approaches can provide an effective way to overcome the complexity of large biological network analysis. Embedding approaches map nodes to low-dimensional vectors by preserving the properties of their higher dimensional counterparts. Lower dimensional embeddings are then used in downstream analysis such as supervised learning link prediction tasks.

Embedding methods have been developed and used in a wide range of domains, most notably natural language processing. While research exploring the application of these methods to biological networks is still in its early stages, there is considerable interest. These applications can be loosely divided into three categories; (1) *drug-related applications*, such as drug-target interactions (DTIs) (4–9), drug-disease interactions (10–12), drug side-effects (13, 14), drug-drug interactions (15–17), polypharmacy antagonistic effects (18, 19) and synergistic reactions in drug combination therapy (20); (2) *protein-related applications*, such as protein-protein interactions (PPIs) (21–24) and protein/gene disease interactions (25–31); and (3) *transcriptomics-related applications*, such as lncRNAs-diseases associations (32–35) and miRNA-disease associations (36–43) and many other applications (44–50).

Since network embedding methods were not originally developed for biological networks, their performance in obtaining different biological network features is yet to be established. Different network embedding approaches capture the network’s structural properties using different methods; the focus can be on local or global properties (51–53). Biological networks are sparse and incomplete (54). Therefore, it is necessary to develop embedding models that take into account the sparsity and incompleteness of biological networks while also accounting for their local and global structural properties. As no single approach appears to handle these trade-offs satisfactorily, one may ask whether integrating network embeddings from different methods might provide richer feature representations, greater insight into the network, and better prediction performance when used in downstream analysis. To address this question we have developed the BioNE processing pipeline, which provides tools for the preparation of networks, application of different network embedding methods and integration of embeddings in different ways. To the best of our knowledge, this pipeline is the first set of tools to support comprehensive integration of network embeddings.

## Implementation

The BioNE pipeline consists of three steps: network preparation, network embedding, and link prediction:

a. Network preparation involves representing the input data in formats suitable for processing within the network embedding step. In this step, users are required to convert adjacency matrices to edge list files. The user also has an option to set networks as directed or two or more edge lists can be combined to create heterogeneous networks.
b. BioNE’s network embedding step takes the prepared input and applies network embedding methods to learn low-dimensional vector representations for each node on the network. The following embedding methods are available within BioNE; LINE (55), GraRep (56), SDNE (57), HOPE (58), LaplacianEigenmaps (LAP) (59), node2vec (60), DeepWalk (61) and Graph Factorization (62). The user has options to treat the network as directed, weighted or set the vector representations size. The output is a space-delimited file that contains vector representations (features/embeddings) of the nodes.
c. In the link prediction step, the user needs to provide the annotation file in order to define dependent and independent variables for link prediction tasks. For example, the annotation file should contain information such as, drug A and protein E do not show an association; drug B and protein E show an association. Therefore, the dependent variable of link drug A and protein E is 0 and for drug B and protein E is 1. The concatenation of the drug and protein embeddings are independent variables (see Figure 1). The user is also required to provide the list of embedding files generated by different embedding methods (produced in the embedding step), which the user wishes to integrate. The user can select the cross-validation method and parameters, an oversampling method (for imbalanced data), feature selection techniques and prediction models.

**Fig. 1.**
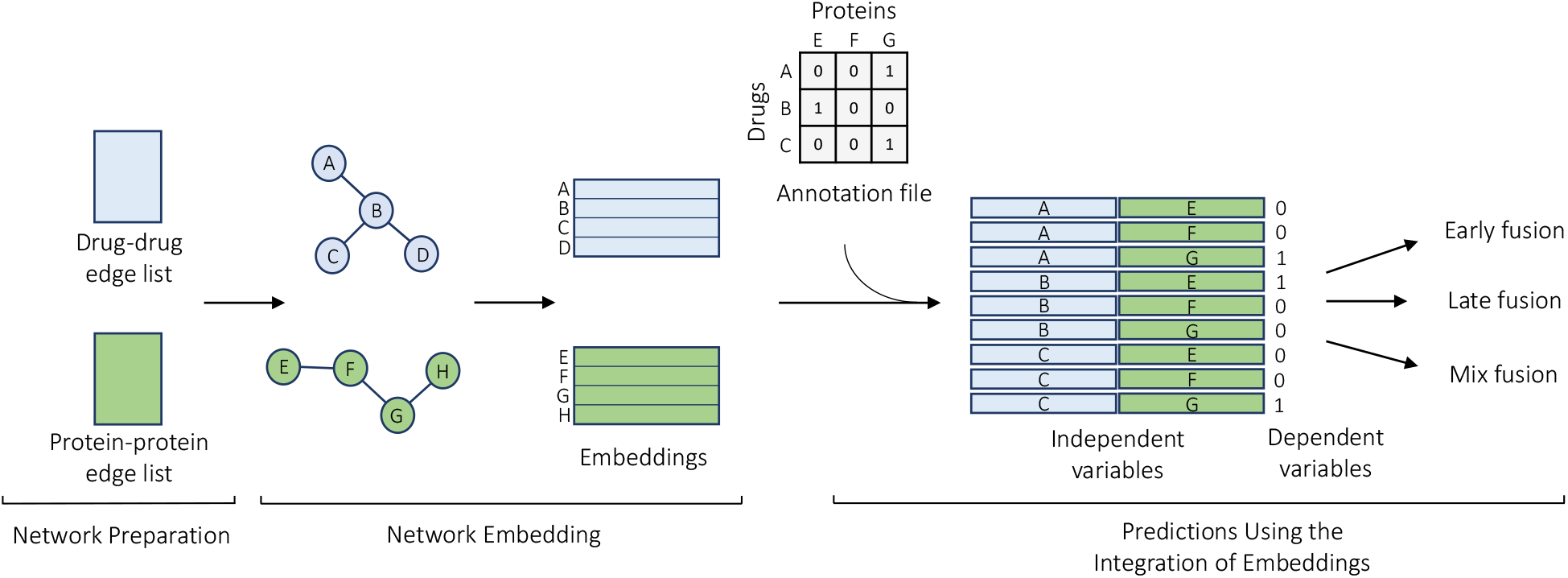
Use case example of BioNE in the use of embedding methods in a drug-target interaction prediction task without the application of the fusion step. This consists of three parts: preparation of the networks, learning of vector representations using network embedding methods, and predictions using the learnt representations. In this prediction model, embeddings are independent variables and their interactions (i.e. 0 and 1) in the annotation file (i.e. known drug-target interactions) are dependent variables.

Three techniques, late (eq.1), early (eq.2) and mixed (eq.3) fusions are used to integrate different embeddings.

### Late Fusion

For late fusion the resulting embeddings are fed to the classifier (machine learning model) separately. The classifier then calculates the prediction probabilities for each instance. The class with the highest sum of prediction probabilities for each instance is assigned as the prediction, that is:

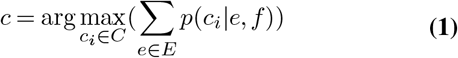

where *C* is the set of classes 0, 1, *E* is the set of embeddings {*e*_*line*_, *e*_*grarep*_, *e*_*sdne*_, *e*_*hope*_, *e*_*lap*_, *e*_*node*2*vec*_, *e*_*deepwalk*_, *e*_*gf*_ } derived from different embedding methods {*line, grarep, sdne, hope, lap, node*2*vec, deepwalk, gf*}, and *p*(*c*_*i*_ | *e, f*) is the prediction probability of class *c* for classifier *f* (e.g. support-vector machine, SVM) for data *e*.

### Early fusion

Early fusion concatenates the embedding results before inserting them into the prediction model. In the case of the integration of grarep, SDNE and deepwalk embeddings:

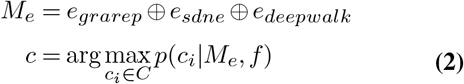

where *M*_*e*_ is a concatenation of embeddings. The classifier *f* (i.e. SVM) then estimates the conditional class probabilities *p*(*c*_*i*_|*M*_*e*_, *f*) for the merged data *M*_*e*_.

### Mixed fusion

Mixed fusion merges data as in early fusion and then passes them on to different classifiers. The sum of prediction probabilities from different classifiers for each instance is calculated and the class with the highest sum is considered as the prediction:

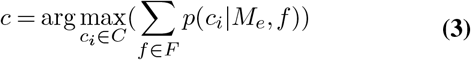

where *M*_*e*_ is a concatenation of embeddings of methods and *F* is the set of classifiers {*Random F orest, SV M, N aive Bayes, XGBoost*} and *p*(*c*_*i*_ | *M*_*e*_, *f*) is the prediction probability of class *c* for classifier *f* for data *M*_*e*_.

BioNE outputs the prediction performance of the link prediction in different metrics. In addition, BioNE also provides receiver operating characteristic (ROC) and precision-recall (PR) curves. ROC shows the true positive rate of a model’s prediction plotted against its false positive rate as the classification threshold varies. The PR curve displays the trade-off between precision and recall for different classification thresholds. Both plots help evaluate the model by plotting performance trade-offs; ROC are the most common of the two, but PR curves can be useful for highly imbalanced classes, which is a common feature in many biological prediction tasks.

### Advantages of BioNE

As mentioned above, the BioNE framework integrates embeddings from different embedding method, enabling the assessment of whether the combined embeddings offer complementary information with regards to the input network features and thus better performance on prediction tasks. In addition, BioNE provides toolsets to overcome the challenges in link prediction and machine learning analysis. In link prediction tasks, machine learning classifiers are used to classify the presence or absence of an interaction between two entities. In biological networks, there is often the problem of class imbalance, as the absence of interactions tend to overwhelmingly dominate the distribution. BioNE provides different oversampling techniques such as SMOTE (63) to overcome this challenge. Another oversampling technique available in BioNE is to equalize the number of positive and negative interactions.

Following the integration of embeddings, the total number of features increases and can lead to the “curse of dimensionality” which can cause substantial issues for most traditional machine learning algorithms. Insufficiency of training samples and redundancy among features is regarded as a significant issue in the supervised classification of hyperspectral data. BioNE provides feature selection methods based on ANOVA and Mutual information (MI) to address this issue. In addition, BioNE provides different machine learning models and helps reduce the risk of over-fitting by providing two different cross-validation methods, namely, k-fold and stratified cross-validation. The performance of the prediction is evaluated using different metrics such as accuracy, precision, recall, specificity, F-scores and area under the ROC and PR curves. As mentioned previously, the PR curve, which mainly focuses on true positive cases, is particularly valuable when measuring model link prediction performance on imbalanced data.

### Example Application of BioNE

As an example of the use of BioNE, the late fusion technique has been tested on the drug-target interaction (DTI) prediction task shown in Figure 1. To conduct this prediction, drugdrug interactions (708 drugs) and known DTIs were extracted from the Drug-Bank database (Version 3.0) (64). Proteinprotein interactions (1493 proteins) were obtained from the Human Protein Reference Database (HPRD, release 9) (65). Embedding methods (LINE, GraRep, SDNE, HOPE, LAP, DeepWalk and GF) were applied to both drug-drug interactions and protein-protein interactions networks separately using default hyperparameters. Detailed explanation of parameters can be found on the Github^1^ page. Vector representations of the drugs are derived from the embedding of drug-drug interaction networks, and vector representations of the proteins are derived from the embeddings of proteinprotein interaction networks. The size of vector representations achieved from each network embedding method is 20. The embeddings of the proteins and drugs are then concatenated according to the annotation file that contains known DTIs. These concatenations are considered as predictors, and the absence or presence of associations between drugs and proteins are taken to be the values of the dependent variables, as in (66).

The BioNE pipeline easily integrates different embeddings achieved from network embedding methods and uses the late fusion technique to test the performance of the predictions. For the late fusion step, a 10-fold cross-validation procedure without the application of oversampling and feature selection methods was conducted. In order to eliminate the problem raised due to imbalanced size between classes, the number negative associations is matched to the number of positive associations, as done in other studies (67). This is achieved by randomly (under)sampling the specified number of negative associations from the sample. In this task, the prediction model used was SVM with a radial basis function (RBF) kernel. In addition, we assessed several embeddings individually for comparison to BioNE’s late fusion. Prediction performance of this method and other embedding methods is shown in Table 1. This table summarizes the performance of this prediction for each fold, reported values include the mean performance metrics. This table, along with ROC and PR curves are the outputs of the BioNE framework.

**Table 1.** Cross validation results for late fusion, as implemented through the BioNE pipeline: different metrics to evaluate the performance of the late fusion in the drugtarget interaction prediction task. Each line represents the performance in each fold of cross-validation and last line takes the mean of these metrics.

|  | Accuracy | Precision | Recall | F1 | ROC-auc | PR-auc |
| --- | --- | --- | --- | --- | --- | --- |
| fold1 | 0.80 | 0.77 | 0.78 | 0.78 | 0.87 | 0.88 |
| fold2 | 0.80 | 0.79 | 0.76 | 0.77 | 0.87 | 0.85 |
| fold3 | 0.72 | 0.75 | 0.70 | 0.73 | 0.83 | 0.86 |
| fold4 | 0.83 | 0.85 | 0.81 | 0.83 | 0.88 | 0.90 |
| fold5 | 0.83 | 0.90 | 0.79 | 0.84 | 0.89 | 0.93 |
| fold6 | 0.82 | 0.86 | 0.78 | 0.82 | 0.91 | 0.93 |
| fold7 | 0.81 | 0.87 | 0.75 | 0.80 | 0.90 | 0.91 |
| fold8 | 0.77 | 0.75 | 0.74 | 0.74 | 0.85 | 0.86 |
| fold9 | 0.77 | 0.83 | 0.70 | 0.76 | 0.85 | 0.86 |
| fold10 | 0.76 | 0.81 | 0.71 | 0.76 | 0.86 | 0.89 |
| Mean | 0.79 | 0.82 | 0.75 | 0.78 | 0.87 | 0.89 |

In addition, Table 2 compares the area under ROC and PR curves of other network embedding methods in the drugtarget interaction prediction task.

**Table 2.**
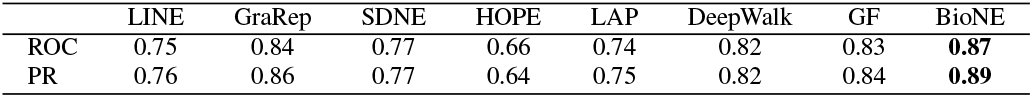
Mean area under the ROC and PR curves in different embedding methods in DTI prediction task. The column labelled BioNE shows the results of predictions using late fusion for combining different embedding methods.

Results show that the BioNE results outperform other network embedding methods when comparing area under ROC and PR curves. With area under the ROC curve of 0.87 BioNE outperformed network embedding methods GraRep, GF, DeepWalk, SDNE, LINE, LAP and HOPE by %3, %4, %5, %10, %12, %13 and %21 respectively. This demonstrates that, BioNE is a valuable tool set to easily integrate the embeddings to achieve more comprehensive knowledge of the network and to test their performance in prediction tasks.

## Conclusions

As network embedding methods were not originally designed for biological networks, their performance in obtaining different biological network features is not straightforward. We believe, the integration of network embeddings can take into account the sparsity and incompleteness of biological networks and is also capable of hurdling the trade-off between local and global structure properties in network embedding methods.

Therefore, the main purpose of this framework is to enable researchers to easily reuse and combine well-known network embedding methods and test their performance individually and in combination on different link/association predictions. In addition, BioNE provides tool sets to overcome some of the challenges in link prediction and machine learning methods such as oversampling methods to overcome the imbalanced data challenges, feature selection methods to reduce the curse of dimensionality and many other tools. Although, we focused on well-known embedding methods, users can expand this framework by adding other de-novo network embedding methods such as graph convolutional networks (GCN) (68, 69), other feature selection and oversampling methods.

In addition, in the case where users wish to add other features unrelated to the networks to the prediction task, they can integrate these features to vector representations (embedding step’s output) and then pass them as inputs to the prediction task. To the best of our knowledge, this pipeline is the first easy to use toolset to support comprehensive integration of network embeddings and test their performance in prediction task, and we intend to develop it further in cooperation with its user community.

## ACKNOWLEDGEMENTS

The author P.P. would like to acknowledge Melisa Chuong for their help in proof reading during the preparation of this manuscript.

## AVAILABILITY OF DATA AND MATERIALS

The datasets supporting the conclusions of this article, as well as BioNE pipeline and detailed explanation of implementation are freely available on GitHub, at https://github.com/pooryaparvizi/BioNE.

## AVAILABILITY AND REQUIREMENTS OF BioNE

Project name: BioNE

Project home page: https://github.com/pooryaparvizi/BioNE

Archived version: https://doi.org/10.5281/zenodo.5500712 Operating system(s): Platform independent

Programming language: Python Other requirements: Bash

License: GNU General Public License v3.0 Any restrictions to use by non-academics: No

## ETHICS APPROVAL AND CONSENT TO PARTICIPATE

No ethical approval or consent to participate required. All of the data used in this study are publicly available.

## CONSENT FOR PUBLICATION

Not applicable.

## AUTHOR CONTRIBUTIONS

P.P. designed, developed, implemented and wrote the manuscript; S.L, F.A. and E.T. supervised P.P. in this study and reviewed the development and implementation of the pipeline and edited the manuscript.

## Footnotes

1 https://github.com/pooryaparvizi/BioNE

